# MLL4 is a critical mediator of differentiation and ferroptosis in the epidermis

**DOI:** 10.1101/2021.05.03.442432

**Authors:** Shaun Egolf, Jonathan Zou, Amy Anderson, Yann Aubert, Kai Ge, John T. Seykora, Brian C. Capell

## Abstract

The epigenetic regulator, *MLL4* (*KMT2D*), has been described as an essential gene in both humans and mice^1,2^. In addition, it is one of the most commonly mutated genes in all of cancer biology^3–7^. Here, we identify a critical role for Mll4 in the promotion of epidermal differentiation and ferroptosis, a key mechanism of tumor suppression^8,9^. Mice lacking epidermal *Mll4*, but not the related enzyme *Mll3* (*Kmt2c*), display features of impaired differentiation and human pre-cancerous neoplasms, including epidermal hyperplasia, atypical keratinocytes, and a loss of polarization, all of which progress with age. Mll4 deficiency profoundly alters epidermal gene expression, and uniquely rewires the expression of key genes and markers of ferroptosis (*Alox12*, *Alox12b*, *Aloxe3*)^10^. Beyond identifying a novel role for *Mll4*-mediated tumor suppression in the skin, our data reveal a potentially much more broad and general role for ferroptosis in the process of epidermal differentiation and skin homeostasis.

---

Epigenetic dysregulation is a “hallmark” of human cancer^11^, with almost half of all human cancers bearing mutations in epigenetic regulators. Squamous cell carcinomas (SCCs) occur on a variety of epithelial surface tissues and harbor the highest rates of mutations in chromatin modifying enzymes among cancers^12^. The epigenetic regulator MLL4 (KMT2D) is one of the most frequently mutated genes across all cancers^6,7^. Notably, mutations in *MLL4* have been associated with both early clone formation in normal epithelial tissues^13–15^, as well as with aggressive and metastatic forms of cutaneous SCC (cSCC)^16–18^, the second most common of all human malignancies^19^. The emerging picture, from both these and other studies^6,7^, suggests that *MLL4* typically accumulates loss of function mutations that impair its ability to suppress both the initiation and progression of cancer.

Despite this, while previous work from our lab has identified a critical role for MLL4 in epidermal gene regulation^20^, the mechanisms behind MLL4-mediated tumor suppression in the skin is virtually unknown. To address this, we created mice with epidermal-specific deletions of *Mll4*, as the total body knockout of *Mll4* is embryonic lethal at day 9.5^2^. We utilized a previously described conditional knockout of *Mll4* in which the catalytic SET domain is flanked by two floxP sites (exons 50 and 51) and whereby Cre-mediated deletion of the floxed catalytic SET domain destabilizes the protein and results in tissue-specific deletion of Mll4^21^. We crossed this genetic model with keratin 14 Cre (Krt14-Cre) expressing mice to generate mice with epidermal deletions of *Mll4* (Krt14-Cre; *Mll4*^fl/fl^ or “Mll4-eKO”) (Extended Data Fig. 1A). Mll4-eKO mice were born in the expected Mendelian ratios (Extended Data Fig. 1B) with no visible differences from littermate controls during the first week of life. However, at approximately 21 days after birth, Mll4-eKO mice displayed a striking cutaneous phenotype marked by visibly red, scaly skin with scattered regions of significant hair thinning (Fig. 1A). In addition to this, Mll4-eKO mice were visibly smaller (Fig. 1A) and weighed 40% less than controls (Extended Data Fig. 1C). This weight difference was consistent across genders (Fig. 1B). At the histological level, Mll4-eKO mice displayed multiple consistent abnormalities in the epidermis as revealed by hematoxylin and eosin (H&E) staining, including overall tissue disorganization and scattered regions of notable epidermal hyperplasia and hyperkeratosis (Fig. 1C). These regions of hyperplasia and hyperkeratosis were notable for both the presence of atypical keratinocytes with large nuclei and increased numbers of mitotic cells (Fig. 1C, inset), all features commonly observed in the precancerous form of cSCC in humans known as actinic keratoses (AKs). As Krt14 is also expressed in the oral and esophageal epithelia, we also examined these tissues. Similar to the epidermis, we observed scattered, albeit more rare regions of epithelial hyperplasia in the oral epithelium of the tongue (Extended Data Fig. 1D), while the esophageal epithelium appeared largely unremarkable (Extended Data Fig. 1E).

**Figure 1.**
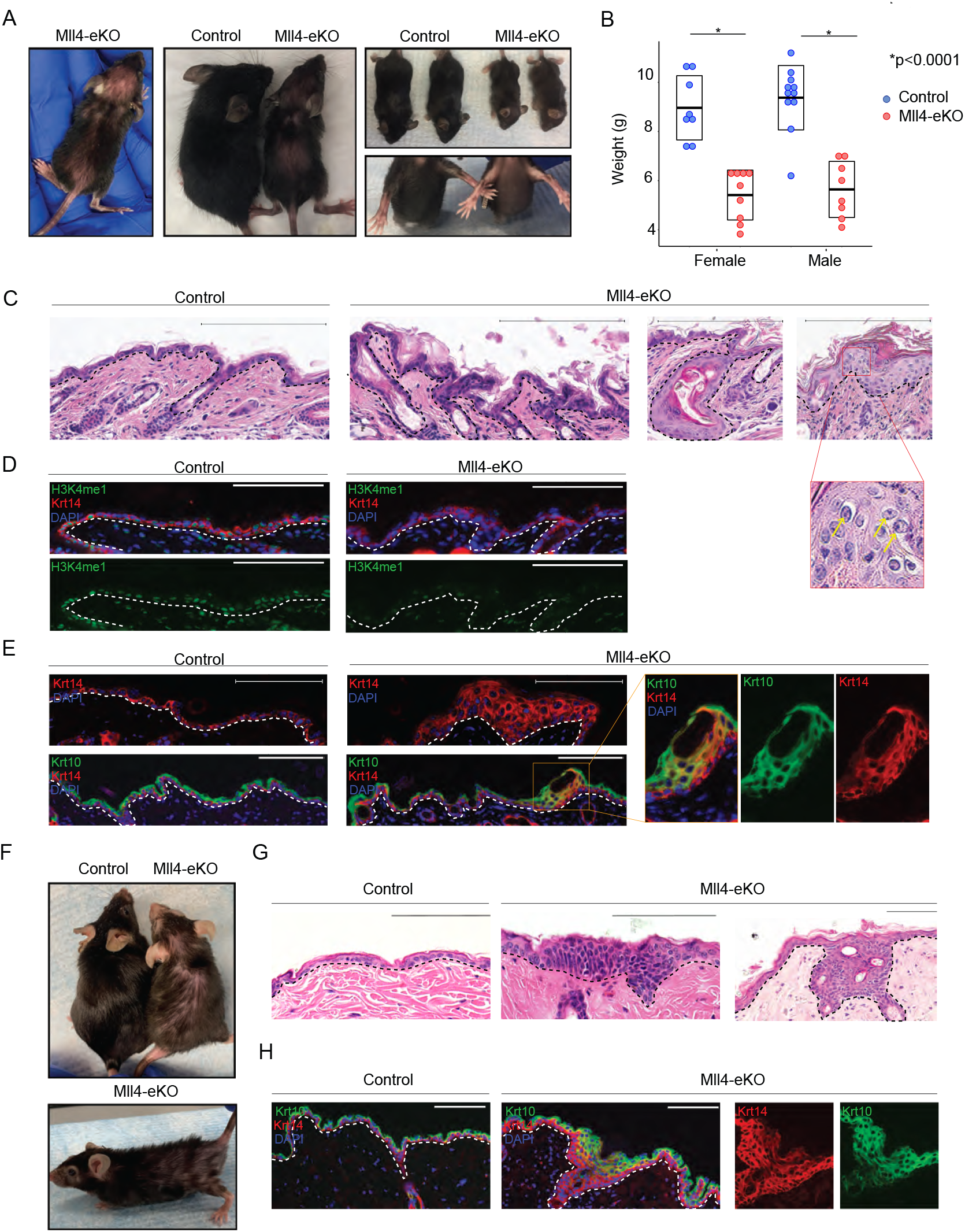
Mll4 deficiency in the epidermis promotes altered differentiation and neoplastic proliferation. **(A)** Representative 3-week-old Mll4-eKO and control mice. **(B)** Weights of either male or female 3-week-old Mll4-eKO (red) and control (blue) mice (n=17-19 mice per genotype, all p<0.0001). **(C)** H&E staining of 3-week-old Mll4-eKO and control mice epidermis (n=17-19 mice per genotype, scale bar: 200 μM). **(D)** IF staining of 3-week-old Mll4-eKO and control mice epidermis for H3K4me1 (green), Keratin-14 (red), and DAPI (blue) (n=5 mice per genotype). **(E)** IF staining of 3-week-old Mll4-eKO and control mice epidermis for Keratin-14 (red), Keratin-10 (green), and DAPI (blue) (n=5 mice per genotype, box denotes zoomed in section). **(F)** Representative 1–2-year-old Mll4-eKO and control mice. **(G)** H&E histological staining 1–2-year-old Mll4-eKO and control mice epidermis (n=3 mice per genotype). **(H)** IF staining of 1–2-year-old Mll4-eKO and control mice epidermis for Keratin-14 (red), Keratin-10 (green), and DAPI (blue) (n=3 mice per genotype). Black and white dashed lines delineate the epidermal and dermal boundary. Scale bar: 100 μM unless otherwise noted.

In the skin, Mll4-eKO mice also presented with sporadic improperly formed hair follicles that resulted in the formation of cyst-like structures (Fig. 1C), consistent with the observed hair thinning and regions of alopecia seen in these mice (Fig. 1A). Given Mll4’s role in catalyzing histone H3 lysine 4 monomethylation (H3K4me1), we examined the global levels of this modification that marks gene enhancers by immunofluorescence (IF) and observed significantly reduced levels (Fig. 1D). These features were not observed in littermate controls which had a histologically normal epidermis.

Given these histological anomalies, we next assessed the various layers of the epidermis which correspond with progressive differentiation states. As expected, control samples had a single layer of keratin 14 (Krt14)-positive staining cells marking the proliferative basal stem cell layer and a distinct, more terminally differentiated keratin 10 (Krt10)-staining spinous layer (Fig. 1E). In contrast, Mll4-eKO mice displayed a multi-layered, expanded Krt14 layer suggestive of a potential failure of upward differentiation of the basal stem cells (Fig. 1E). Further, the expanded Krt14 layers in the epidermis of Mll4-eKO mice commonly contained dual Krt14- and Krt10-positive staining keratinocytes further suggesting a state of dysfunctional epidermal differentiation (Fig. 1E). As increased inflammation from barrier disruption can drive epidermal hyperplasia in different contexts^22^, we investigated levels of inflammation utilizing both H&E and CD3 T cell staining. In both cases, Mll4-eKO did not demonstrate any difference in the amount or numbers of immune cells (Extended Data Fig. 1F, G), suggesting that the significant hyperplasia we observed was not just secondary to increased inflammation.

To observe how Mll4 loss impacts epidermal homeostasis and differentiation over time, we aged Mll4-eKO mice for at least one year and some for up to two years. The epidermal phenotype persisted in Mll4-eKO and appeared to worsen with age, including progression of both the grossly visible hair loss and scaly, hyperkeratotic skin (Fig. 1F). Similarly, at the histological level, the abnormal epidermal phenotype seemed to progress over time, as Mll4-eKO mice displayed regions of neoplastic proliferation, many of which appeared to be more infiltrative and consistent with early cSCC lesions in humans (Fig. 1G). Consistent with this observation, an expanded Krt14/Krt10 dual-staining layer was also observed in these mice (Fig. 1H). Together, these data demonstrate that Mll4 loss significantly impairs proper epidermal homeostasis and differentiation *in vivo*.

To begin to understand the mechanisms behind these changes, we performed RNA-seq on the epidermis from littermate control and Mll4-eKO mice. As expected, given Mll4’s critical role in transcriptional regulation, Mll4-eKO mice demonstrated substantial transcriptional alterations with roughly equal number of significantly downregulated and upregulated genes (Fig 2A, Extended Data Table 1). Gene ontology (GO) analysis of downregulated genes demonstrated a significant enrichment of genes involved in lipoxygenase activity and lipid metabolism (Fig. 2B). This was particularly compelling given the essential role of both of these processes in normal epidermal homeostasis and barrier formation^23,24^. Underscoring the critical nature of many of these downregulated genes, mutations in a number of them are causative in a variety of human genetic skin disorders such as *Col7a1*, *Fermt1*, *Slurp1*, *Alox12b* and *Aloxe3*^25–27^.

**Figure 2.**
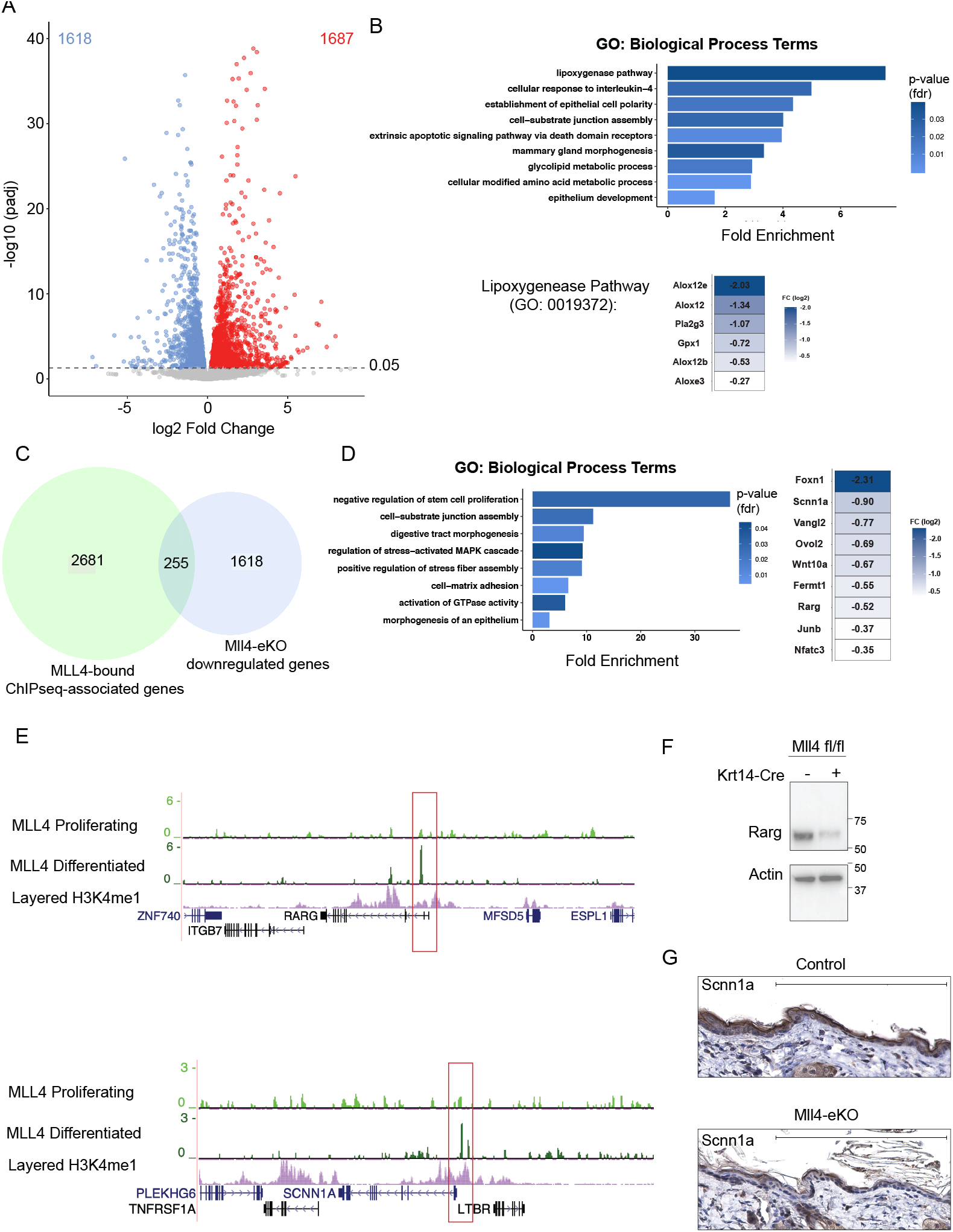
Loss of Mll4 leads to a profoundly altered transcriptome characterized by a loss of key genes involved in differentiation and lipid metabolism. **(A)** Differentially expressed genes (1618 downregulated genes, 1687 upregulated genes) in isolated bulk epidermis of 3-week-old Mll4-eKO and control mice. Dashed line denotes adjusted p-value of 0.05 (n= 3-4 mice per genotype). **(B)** GO biological process analysis of Mll4-eKO differentially expressed downregulated genes (1618 genes). Downregulated genes driving the top GO term “lipoxygenase pathway” are highlighted. **(C)** Intersection of all MLL4-bound region-associated genes in NHEKs and downregulated genes in Mll4-eKO mice (255 genes). **(D)** Left: GO biological process analysis of the intersection in Fig. 2D (255 genes). Right: List of key epidermal differentiation genes enriched in GO terms from Fig 2D intersection. **(E)** ChIP-seq for MLL4 in proliferating or differentiated primary human keratinocytes at key epidermal differentiation (*RARG*) and barrier (*SCNN1A*) genes listed in Fig 2E (n=2-3 per condition). **(F)** Immunoblot of Rarg and Actin isolated from bulk epidermis of 3-week-old Mll4-eKO and control mice. Molecular ladder shown to the right of blot (n=2 mice per genotype). **(G)** IHC staining of Sccn1a of 3-week-old Mll4-eKO and control mice epidermis (n=3 mice per genotype, scale bar: 200 μM).

In contrast, genes upregulated in the Mll4-eKO mice were enriched for numerous genes and GO terms involved in various aspects of fatty acid metabolism, such as fatty acid synthesis, beta-oxidation, and elongation (Extended Data Fig. 2A, Table 1). This was intriguing given that fatty acid metabolism has been implicated in promoting both metastasis and treatment resistance in cancer^28–31^. Notably, many of these significantly upregulated genes have been reported to be involved in driving cancer invasion and metastasis such as *Acat* (*Soat1*), *Cd36*, *Fabp5*, *Mgll*, *Scd1*, *Slc1a5*, and *Slc7a11*^32,33^. For example, CD36 has been shown to mark metastasis-initiating cells in oral SCC and targeting it inhibits metastasis *in vivo*^34^.

To examine whether Mll4 might directly regulate these genes, we differentiated primary normal human epidermal keratinocytes (NHEKs) *in vitro* and mapped Mll4 binding by chromatin immunoprecipitation-sequencing (ChIP-seq). Intersection of Mll4 peaks with genes significantly downregulated in Mll4-eKO mice demonstrated substantial overlap (Fig. 2C). These again included numerous genes involved in epidermal development, differentiation, and barrier formation such as the retinoic acid receptor gene, *RARG* and *SCNN1A* (Fig. 2D, E)^35,36^. Notably, the ChIP-seq data demonstrated that the MLL4 peaks emerged at many of these genes only in the differentiated NHEKs and were not present in the proliferating, undifferentiated NHEKs (Fig. 2E). Specifically, there were 1,389 peaks that were unique to the differentiated state, while only 209 were unique to the proliferating stem cell state (Extended Data 2B, C, Table 2). The majority of peaks (1,870) were common to both (Extended Data 2B, C, Table 2). Consistent with previous data^20^, motif analysis demonstrated that the top transcription factor motifs for MLL4 binding enrichment were those for the master epithelial transcription factor, p63, and p53 (Extended Data Fig. 2D). Western blotting and immunohistochemistry (IHC) confirmed that these gene expression changes resulted in reduced protein levels for RARγ and SCNN1A, respectively (Fig. 2F, G). Collectively, these data demonstrate that mice lacking Mll4 in the epidermis display broad gene expression alterations marked by the loss of key genes and pathways involved in epidermal differentiation and barrier formation, and consistent with the significant phenotype observed in Mll4-eKO mice. Furthermore, MLL4 binds directly at enhancers near many of these genes during the course of differentiation, indicating a potentially direct role in activating their expression.

Like *MLL4*, the related histone methyltransferase, *MLL3* (*KMT2C*), also catalyzes H3K4me1and is frequently mutated in numerous human cancers, including SCCs^3–7^. Therefore, we also wanted to investigate the role of MLL3 in epidermal biology and tumor suppression. To do this, we took a similar approach to generate mice with epidermal-specific deletions of *Mll3* by crossing Krt14-Cre mice with mice carrying *Mll3* alleles with loxP sites flanking a 61 amino acid region of the catalytic SET domain. Cre-mediated recombination results in deletion of the targeted region (Extended Data 3A) and results in destabilization and loss of the full protein^37,38^. In clear contrast to Mll4-eKO mice, mice lacking Mll3 in the epidermis (“Mll3-eKO”) did not display any obvious phenotypic alterations (Fig. 3A) beyond subtle changes around the eye that only appeared with aging (Extended Data 3B). There was no significant change in weight between control and Mll3-eKO mice (Fig. 3B). Histologically, the Mll3-eKO mice also did not demonstrate significant or consistent changes in epidermal thickness or organization (Fig. 3C), and H3K4me1 levels were only minimally reduced (Fig. 3D). Consistent with these findings, Krt14 and Krt10 staining did not suggest alterations in differentiation dynamics in contrast to Mll4-eKO mice (Fig. 3E). Finally, RNA-seq of Mll3-eKO mice showed that a loss of *Mll3* in the epidermis only led to modest alterations in gene expression (Fig. 3F, Extended Data Table 3), with minimal overlap with those genes altered by Mll4-deficiency (Extended Data Fig. 3C). Together, these data demonstrated that Mll4 plays a more critical role in epidermal gene regulation, homeostasis and differentiation than Mll3.

**Figure 3.**
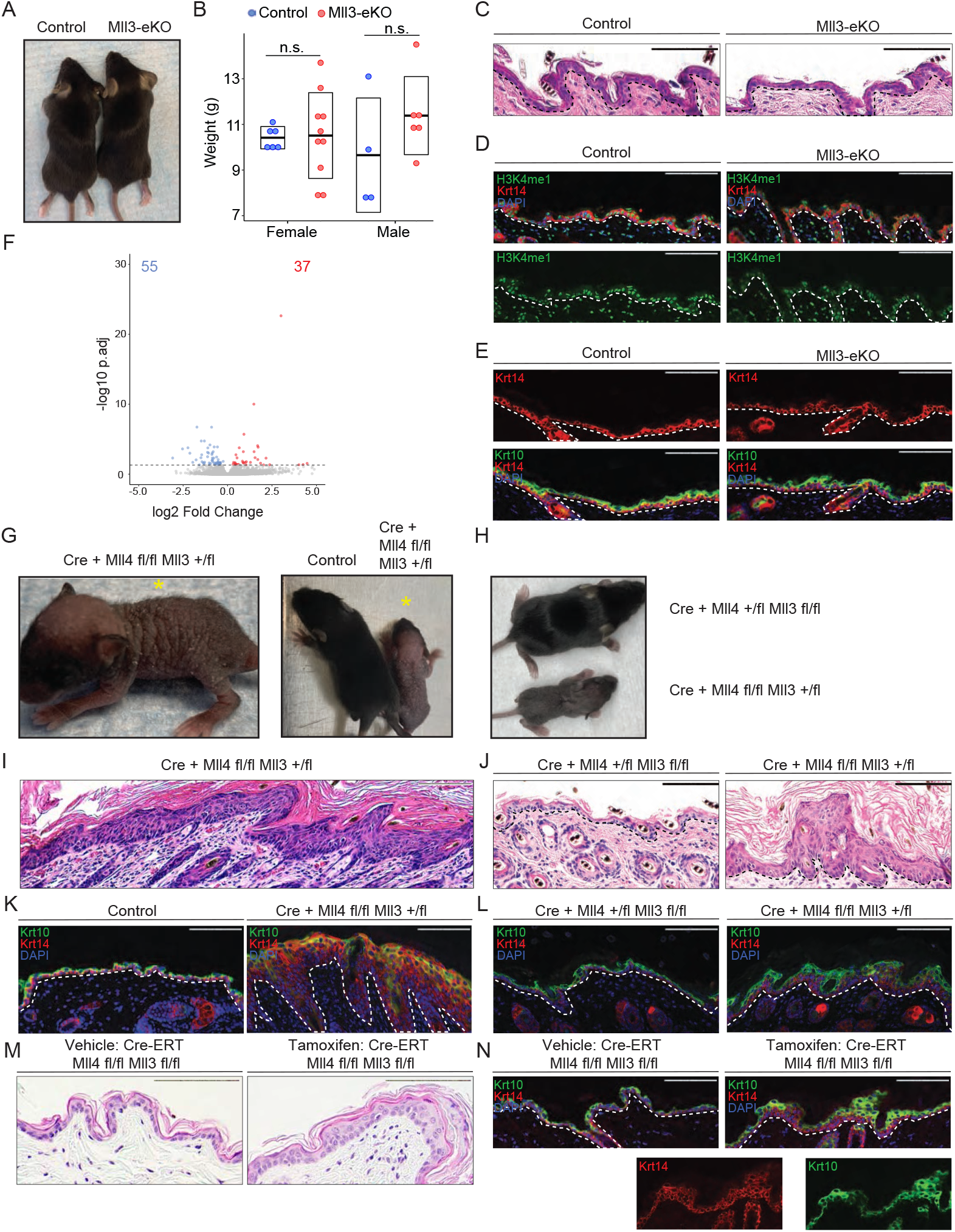
In contrast to Mll4 loss, Mll3 deficiency leads to modest effects in the epidermis. **(A)** Representative 3-week-old Mll3-eKO and control mice. **(B)** Weights of either male or female 3-week-old Mll3-eKO (red) and control (blue) mice (n=9-16 mice per genotype, p=0.49). **(C-E)** Comparison of 3-week-old Mll3-eKO and control mice epidermis by **(C)** H&E staining, **(D)** IF staining of H3K4me1 (green), Keratin-14 (red), and DAPI (blue) or **(E)** IF staining of Keratin-14 (red), Keratin-10 (green), and DAPI (blue) (n=3 mice per genotype). **(F)** Differentially expressed genes (55 downregulated genes, 37 upregulated genes) in isolated bulk epidermis of Mll3-eKO and control mice. Dashed line denotes adjusted p-value of 0.05 (n= 6 mice per genotype). **(G)** Representative 1-week-old Krt14-Cre(+); *Mll4*^fl/fl^; *Mll3*^+/fl^ mouse alone (left) or beside a littermate control (right). **(H)** Littermate sibling Krt14-Cre(+); *Mll4*^fl/+^; *Mll3*^fl/fl^ (top) and Krt14-Cre; *Mll4*^fl/fl^; *Mll3*^+/fl^ (bottom) mice compared. **(I)** H&E staining of a Krt14-Cre(+); *Mll4*^fl/fl^; *Mll3*^+/fl^ mouse epidermis (n=3 mice per genotype). **(J)** Littermate sibling Krt14-Cre(+); *Mll4*^fl/+^; *Mll3*^fl/fl^ and Krt14-Cre(+); *Mll4*^fl/fl^; *Mll3*^+/fl^ mice compared by **(J)** H&E histological staining of the epidermis, and **(K)** IF staining for Keratin-14 (red), Keratin-10 (green), and DAPI (blue) (n=3 mice per genotype). **(M-N)** Comparison of the epidermis of vehicle and tamoxifen treated Krt14-CreER^tam^; *Mll4*^fl/fl^; *Mll3*^fl/fl^ mice by **(M)** H&E staining or **(N)** IF staining of Keratin-14 (red), Keratin-10 (green), and DAPI (blue) (n=2 mice per group, one vehicle control, one genotype control). Black and white dashed lines delineate the epidermal and dermal boundary. Scale bar: 100 μM unless otherwise noted.

To determine whether Mll3 may compensate on some level for the loss of Mll4 in the epidermis, or whether it was completely dispensable even in the setting of Mll4 deficiency, we attempted to generate mice lacking both Mll3 and Mll4 in the epidermis. Here we observed that mice lacking Mll4 and one copy of Mll3 were even more severely affected than mice lacking just Mll4, suggesting that Mll3 does in fact perform some compensatory functions for Mll4 in its absence. These Krt14-Cre(+); *Mll4*^fl/fl^; *Mll3*^+/fl^ mice displayed markedly wrinkled, scaly skin with sparse hair (Fig. 3G). Notably, these mice were significantly more affected than mice lacking all Mll3 and one copy of Mll4 (Krt14-Cre(+); *Mll4*^fl/+^; *Mll3*^fl/fl^) (Fig. 3H). At the histological level, Krt14-Cre(+); *Mll4*^fl/fl^; *Mll3*^+/fl^ mice presented with an even more thickened epidermis and expanded epidermis than mice lacking only Mll4 (Mll4-eKO) (Fig. 3I, Extended Data Fig. 3D). In contrast, Krt14-Cre(+); *Mll4*^fl/+^; *Mll3*^fl/fl^ mice (Extended Data Fig. E, F, G) appeared less affected than both Krt14-Cre(+); *Mll4*^fl/fl^; *Mll3*^+/fl^ mice as well as Mll4-eKO mice (Fig. 3K, J, L). Importantly, we found that we were unable to generate any live mice lacking both Mll3 and Mll4, suggesting that at least one copy of these two related enzymes is required for epidermal development (Extended Data Fig. 3H). To test this further, we created a tamoxifen-inducible line of these mice using a Krt14-CreER^tam^ system. Utilizing this approach, we demonstrated that deletion of both *Mll3* and *Mll4* in the adult epidermis recapitulated several of the phenotypic manifestations of embryonic loss of Mll4 and/or Mll3 (Fig. 3M, N). In contrast to the constitutive model, these data demonstrated that complete deletion of both enzymes in the adult epidermis was compatible with life. Collectively, these observations supported the conclusions that Mll4 is indeed the more critical H3K4 monomethylase in the epidermis, and that at least one copy of *Mll3* or *Mll4* is required for epidermal development and viability.

To begin to explain the more severe phenotype observed with Mll4 loss in comparison to Mll3, we took advantage of the significantly different transcriptional profiles that these mice displayed, and the fact that the changes appeared to reflect the observed unique phenotypic manifestations. In particular, the most notable difference observed in the Mll4-eKO mice was the significant loss of expression of key genes involved in lipoxygenase activity such as *Alox12*, *Alox12b*, and *Aloxe3* (Fig. 2D). While lipoxygenases are well-established as key enzymes during the process of epidermal differentiation^24^, we were particularly intrigued by their emerging role in the relatively recently discovered form of programmed tumor suppressive cell death known as ferroptosis that is characterized by lipid peroxidation (Fig. 4A)^8,10,39^. For example, Alox12 is required for p53-mediated tumor suppression via ferroptosis^10^. Many human cancers are noted to have a loss of one copy of *ALOX12* on chromosome 17p^40,41^, and variants in *ALOX12* have also been associated with human SCCs^42^. In addition to the loss of these key lipoxygenase enzymes, two of the key genes which suppress ferroptosis through the elimination of lipid peroxides, *Slc7a11* and *Gpx4*^8–10,43^, were concomitantly significantly upregulated in the Mll4-eKO mice (Fig. 4A). Furthermore, among the upregulated fatty acid metabolism genes we identified in the Mll4-eKO mice (Extended Data Fig. 2A), the gene stearoyl CoA desaturase-1 (*Scd1*), the rate-limiting enzyme monounsaturated fatty acid synthesis, has been shown to both promote cancerous invasion and protect against ferroptosis^44,45^.

**Figure 4.**
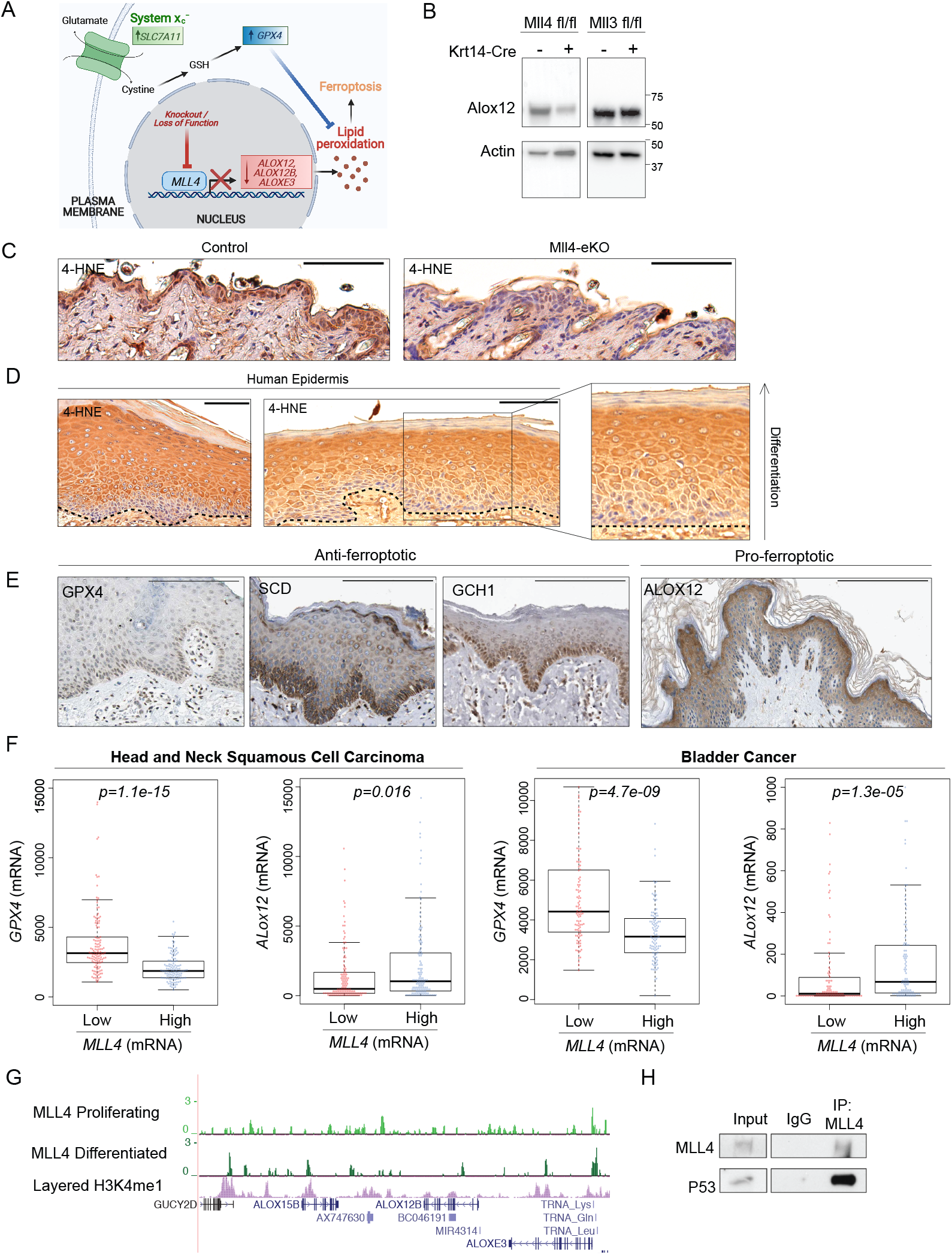
Mll4 deficiency impairs ferroptosis via rewired gene expression. **(A)** Schematic demonstrating that knockout of *Mll4* results in lost expression of pro-ferroptosis genes (*Alox12*, *Alox12b*, *Aloxe3*) and upregulation of key anti-ferroptosis genes (*Slc7a11*, *Gpx4*) ultimately decreasing cellular lipid peroxides and inhibiting ferroptosis. **(B)** Immunoblot of Alox12 and Actin isolated from bulk epidermis of 3-week-old Mll4-eKO, Mll3-eKO, and control mice. Molecular ladder shown to the right of blot (n=2 mice per genotype). **(C)** IHC staining of 4-HNE in 3-8-week-old Mll4-eKO and control mice epidermis (n=5 mice per genotype). **(D)** IHC staining of 4-HNE in human epidermis (n=4 patient samples). Dashed line indicates epidermal and dermal boundary. **(E)** IHC staining of anti-ferroptosis (GPX4, SCD, GCH1) or pro-ferroptosis (ALOX12) proteins in human epidermis obtained from the Human Protein Atlas. Scale bar: 200 μM. **(F)** Comparison of GPX4 and ALOX12 expression in the lowest one-fourth and highest one-fourth of MLL4 expressing tumors in 515 human head and neck squamous cell carcinomas (left) or 411 human bladder cancers (right) obtained from The Cancer Genome Atlas (TCGA). All p<0.05. **(G)** ChIP-seq for MLL4 in proliferating or calcium differentiated human keratinocytes at the Alox family genes (n=2-3 per condition). **(H)** Co-IP of MLL4 followed by immunoblotting of p53 in human keratinocytes (n=3). Scale bar: 100 μM unless otherwise noted.

Given these changes, we first verified Alox12 loss at the protein level in Mll4-eKO, but not Mll3-eKO mice (Fig. 4B). We then utilized a marker of lipid peroxidation and ferroptosis, 4-hydroxynoneal (4-HNE)^8,46^, to assess the impact that loss of Mll4 might have. Strikingly, we observed a marked reduction in 4-HNE staining in Mll4-eKO mice (Fig. 4C) that was not observed in Mll3-eKO mice, suggesting that a potentially significant reduction in Alox-mediated ferroptosis might occur in the setting of Mll4 deficiency. More speculatively, these data provoked the hypothesis that ferroptosis may actually play a critical role in the promotion of epidermal differentiation. Indeed, the precise mechanisms by which keratinocytes enucleate and undergo cell death during terminal differentiation is poorly understood. Therefore, we stained human skin samples to assess how the ferroptosis marker, 4-HNE, may appear. Remarkably, we observed a clear progressive increase in staining intensity as cells progressed towards terminal differentiation, with the highest levels of staining observed in the highly metabolically active stratum granulosum (Fig. 4D), precisely where keratinocytes make the transition to becoming enucleated and forming the stratum corneum where they ultimately slough off the skin.

In support of these observations, data from the Human Protein Atlas^47^ confirmed the enrichment of anti-ferroptotic regulators like GPX4, SCD1, and GCH1 in the proliferative basal stem cell layer, while pro-ferroptotic proteins like Alox12 were enriched in the stratum granulosum where the highest levels of 4-HNE were observed (Fig. 4E). Additionally, other publicly available data from the The Cancer Genome Atlas (TCGA)^3,4^, showed that lower expression of MLL4 was significantly associated with both lower expression of ALOX12 as well as higher expression of GPX4 in human head and neck SCC as well as bladder cancer (Fig. 4F). In support of a direct role for MLL4 in promoting the expression of these lipoxygenase genes, ChIP-seq demonstrated that MLL4 binds at potential regulatory elements and enhancers adjacent to these Alox-family genes (Fig. 4G). Furthermore, in addition to our previous data showing that MLL4 interacts with the transcription factor, p63^20,48^, we identified that MLL4 also interacts with p53 (Fig. 4H). Both p53 and p63 have been shown to induce the expression of Alox12 to promote both epidermal differentiation^49^ and tumor suppression^10,50^, respectively. This is also consistent with our ChIP-seq motif analysis showing that the main transcription factor binding motifs of MLL4 are p63/p53 sites (Extended Data Fig. 2D). Together, these data support a model whereby Mll4 is critical for the activation of key lipoxygenase genes such as *Alox12* in order to promote both epidermal differentiation and barrier formation, and in turn, tumor suppression via ferroptosis (Fig. 4A). Beyond this direct regulation of lipoxygenase genes, the aberrant upregulation of ferroptosis-suppressing genes such as *Gpx4*, *Slc7a11*, and *Scd1* in Mll4-eKO mice supports a potentially lipogenic and ferroptosis-resistant state that confers a cellular advantage to promote the neoplasms observed in these mice.

In summary, our results offer unique insight into the primary importance of MLL4 as compared to MLL3 in the skin epidermis. As *MLL4* (*KMT2D*) loss of function mutations are amongst the most common events in human cancer, the result presented here offer a novel mechanism by which these mutations may promote both the initiation and progression of cancer, particularly in the setting of keratinocyte cancers, which collectively outnumber all other human malignancies^19^. Given the ability to pharmacologically induce ferroptosis through inhibitors of targets such as GPX4 or SLC7A11^8,9^, these data suggest a potentially new therapeutic strategy for treating epithelial cancers like SCC, particularly in the setting of *MLL4* mutations. In addition, in the context of recent evidence demonstrating that drugs which induce ferroptosis can also operate synergistically with immunotherapies^8,51^, their may exist even more potential uses when employed in combination. More generally, as these results implicate ferroptosis in the general process of epidermal differentiation and skin homeostasis, it suggests that future studies may uncover a role for dysregulated ferroptosis not only in skin cancers, but also potentially in a variety of diverse skin disorders as well.

**Supplementary Information** is available for this paper:

**Extended Data Table 1. MLL4-eKO RNA-seq**

**Extended Data Table 2. MLL4 ChIP-seq**

**Extended Data Table 3. MLL3-eKO RNA-seq**

## METHODS

### Normal Human Epidermal Keratinocyte Isolation and Culture

Primary epidermal progenitors were isolated from de-identified discarded neonatal human foreskin obtained by Core B of the Penn Skin Biology and Diseases and Resource-based Center. Foreskin was incubated at 4°C for 12 h in 2.4 U/mL Dispase II. Sterile forceps were used to separate the underlying dermis. The epidermal sheet was transferred to a 60-mm tissue culture plate, incubated in 0.25% trypsin for 10 min at 37°C, and then neutralized with 1 mL of fetal bovine serum (FBS). Sterile forceps were used to scrape the epidermal sheet against the dish to dissociate cells. The suspension was passed through a 40-μm strainer and then centrifuged at 200 g for 5 min. The cell pellet was resuspended in 5 mL keratinocyte medium (described next). Epidermal progenitors were cultured in a 50:50 mix of 1 × keratinocyte–SFM supplemented with human recombinant epidermal growth factor and bovine pituitary extract combined with medium 154 supplemented with human keratinocyte growth supplement and 1% 10,000 U/mL penicillin–streptomycin at 37°.

### Murine Models

All animal protocols were reviewed and approved by the Institutional Animal Care and Use Committee of the University of Pennsylvania. Mice were maintained on a mixed B6/129 background on a standard light-dark cycle. Mice carrying Mll4-SET floxed alleles^21^, Mll3-SET floxed alleles^37^, or a combination of both of these were crossed with Keratin-14-Cre transgenic mice. Krt14-Cre(+); *Mll4*^fl/fl^ (Mll4-eKO), Krt14-Cre(+); *Mll3*^fl/fl^ (Mll3-eKO), “triple-floxed” combinations, i.e. Krt14-Cre(+); *Mll4*^fl/fl^; *Mll3*^fl/+^ or Krt14-Cre(+); *Mll4*^fl/+^; *Mll3*^fl/fl^, or a “quadruple-floxed” combination (Krt14-Cre(+); *Mll4*^fl/fl^; *Mll3*^fl/fl^) were considered mutants. Unless noted otherwise, littermates homozygous for the Mll alleles of interest lacking Krt14-Cre were used as controls. A similar approach was utilized using inducible Krt14-CreER^tam^ (005107, Jackson Labs) transgenic mice. To activate inducible knockouts, tamoxifen (T5648-1G, Sigma) was resuspended in corn oil (C8267-Sigma) and used for injections. Both genotype controls and corn oil (vehicle) controls were used to account for the potential effects of tamoxifen. For genotyping, PCR was done using the Thermo Phire Animal Tissue PCR kit (F140WH) using the following primers: Mll3 (F: GTCATCGGTGTGGTCTGAATGA and R: AACCGGAAGGAGAAGCTTTATGA), Mll4 (F: CAGTTGAGCTAGTCAAGTGATT and R: TTCAATGTGGAGGGGAGTGACAG) and Cre (F: GAACCTGATGGACATGG and R: AGTGCGTTCGAACGCTAGAGCCTGT). Unless otherwise stated, all experimental mice were a mix of male and female. The number of animals used per experiment are stated in the figure legends.

### Murine RNA and Protein Extraction

Murine epidermis was dissociated from the dermis before isolation of bulk RNA and protein. Timepoints were selected as to not include the anagen phase of the hair cycle which compromises the dissociation process. Briefly, following euthanasia of 21-day-old mice, the skin was dissected and the underlying fat pad removed using a scalpel. The resulting tissue was floated dermis side down in 5 U/mL Dispase (Corning) in PBS for 40 minutes at 37°C. The epidermis was then removed using a scalpel and for RNA was flash frozen in Trizol and stored at −80°C until RNA isolation. RNA was extracted using RNeasy kit (Qiagen. #74104) at the same time and date for all mice belonging to a single experimental cohort (ie. RNA-seq) regardless of the date of murine euthanasia to reduce batch effects and stored at −80°C. For protein, epidermis was placed directly into cold PBS and centrifuged at 4°C for 5 minutes at 2500 rpm. To the resulting pellet, protein lysis buffer (Cell Signaling) containing a protease inhibitor cocktail was added and the mixture homogenized, sonicated, rotated at 4°C for 10 minutes, then centrifuged at 4°C at full speed for 10 minutes. Lysates were quantified using the Bradford assay. Frozen lysates were stored at −80°C.

### RT-PCR

cDNA was obtained using High-Capacity RNA-to-DNA kit (ThermoFisher. #4368814). For quantitative real-time PCR, Power SYBR Green PCR Master Mix (ThermoFisher. #4367659) was used. Primer sequences are available on request. RT-qPCR data analysis was performed by first obtaining the normalized QT values (normalized to GAPDH and 18S ribosomal RNA) and the 2−ΔΔCt method was applied to calculate the relative gene expression. ViiA 7 Real-Time PCR System was used to perform the reaction (Applied Biosystems). The average and standard deviations were assessed for significance using a student’s t-test. All p values are noted in figure legends and were considered significant if p<0.05 and non-significant (NS) if p>0.05.

### RNA-sequencing

RNA-seq libraries were prepared at the same time for all samples belonging to a single experimental cohort to reduce batch effects. All RNA-seq libraries were prepared using the NEBNext poly(A) mRNA magnetic isolation module followed by NEBNext Ultra Directional RNA library preparation kit for Illumina. Library quality was checked by Agilent BioAnalyzer 2100 and libraries were quantified using the Library Quant Kit for Illumina. Libraries were then sequenced using a NextSeq500 platform (75-base-pair (bp) single-end reads). All RNA-seq was aligned using RNA STAR^52^ under default settings to Mus musculus GRCm38 FPKM (fragments per kilobase per million mapped fragments) generation and differential expression analysis were performed using DESeq2^53^. Statistical significance was obtained using an adjusted p value (padj) generated by DESeq2 of less than 0.05.

### ChIP-sequencing

ChIP-seq was performed as described previously^54^. Briefly, keratinocytes cultured in 10-cm2 dishes were fixed in 1% formaldehyde for 5 min, and fixation was quenched with the addition of glycine to 125 mM for an additional 5 min. Cells were harvested by scraping from plates and washed twice in 1 × PBS before storage at −80°C. ChIP extracts were sonicated for 15 minutes in a Covaris sonicator. All ChIPs were performed using 500 μg of extract and 2 μg of antibody per sample (anti-KMT2D/MLL4, Sigma # HPA035977). Thirty microliters of Protein G Dynabeads was used per ChIP. ChIP DNA was also used to make sequencing libraries using NEBNext Ultra DNA library preparation kit for Illumina. Library quality was checked by Agilent BioAnalyzer 2100 and libraries were quantified using the Library Quant Kit for Illumina. Libraries were then sequenced using a NextSeq500 platform (75-bp, single-end reads). After sequencing, all data were demultiplexed from the raw reads using Illumina’s BCL2FASTQ from BaseSpace. Further ChIP-seq analysis described below.

### ChIP-sequencing Data Processing

ChIP-seq data was analyzed as previously described^54^. Briefly, FASTQ reads from lanes 1 to 4 were combined for each sample and aligned to *homo sapiens* genome (hg19 UCSC) using bowtie2^55^ (version 2.1) allowing one mismatch per seed (-N 1) and reporting one alignment per read (-k 1). Alignment files were filtered with samtools^56^ (version 1.1) to remove unmapped reads (-F 4) and reads with a mapping quality inferior to 10 (-q 10). After sorting and indexing, alignment files were further filtered to remove reads mapped to mitochondrial chromosome or unplaced contigs (chrM, chrUn, chrN_random). MLL4 (KMT2D) peaks were called with HOMER^57^ (version 4.6). Tag directories were generated for each replicate using an estimated fragment length of 150 base-pair (bp), allowing one tag per bp (makeTagDirectory -fragLength 150 -tbp 1). Tag directories from replicates were combined with HOMER makeTagDirectory using an estimated fragment length of 150 base-pair (-fragLength 150). MLL4 peaks were called for each condition (proliferating or differentiated NHEK) from merged tag directories using their corresponding input with HOMER findPeaks (-style factor -size 200 -fragLength auto -F 2 -fdr 0.005), yielding 967 MLL4 peaks in proliferating NHEKs and 2954 MLL4 peaks in differentiated NHEKs. MLL4 peaks from both conditions were combined, sorted, merged and filtered to remove peaks with less than 1 read per million (RPM) yielding 3468 peaks and condition-selective MLL4 peaks were defined by a fold change threshold of 2, yielding 1870 common peaks, 209 proliferation-selective MLL4 peaks and 1369 differentiation-selective MLL4 peaks. Analysis of transcription factor binding motifs at each peak set was performed using HOMER findMotifsGenome.pl script with default parameters, scaling sequence logos by information content (-bits).

### Gene Ontology Analyses

All GO analyses were performed using PANTHER at http://pantherdb.org/ to determine statistically over-represented GO terms using Fisher’s Exact test under the category “Biological Process”. P-values for GO terms are FDR statistics. The top 8 plotted GO terms represent the GO terms with the highest fold enrichment under PANTHER’s default hierarchical clustering categorization. GO term figures generated using ggplot2.

### Co-immunoprecipitation experiments

Co-immunoprecipitations (Co-IP) experiments were performed as previously described^58^. Briefly, 30 mL of magnetic Protein G Dynabeads were washed twice in 1 mL BSA 0.5%, resuspended in 250 mL BSA 0.5% and conjugated for 1h to 2h at 4°C under rotation with 1 mg of antibodies against either MLL4 or IgG as a negative control. About 500,000 proliferating primary NHEKs were harvested from a 10 cm culture plate at 50% confluence and lysed for 1h at 4°C under rotation in 250 mL of immunoprecipitation buffer (IP buffer: 20 mM Tris pH 7.5, 134 mM NaCl, 1 mM CaCl2, 1% NP-40, 10% 36 glycerol, supplemented with freshly made 1 mM MgCl2, 1:100 Halt protease and phosphatase inhibitor cocktail (Thermo Fisher, Cat# 78440) and benzonase at 12.5 U ml-1. Benzonase is critical for the efficient release of chromatin-bound proteins to the supernatant and MgCl2 is critical for its activity. Cell lysates were centrifuged at top speed for 10 minutes. 500 mg cell lysate were incubated overnight at 4°C with antibody conjugated magnetic beads previously washed 3 times with 1mL BSA 0.5%. Immunoprecipitates were collected using a magnet, washed four to five times with IP buffer devoid of MgCl2, protease/phosphatase inhibitors and benzonase, then boiled with NuPage loading dye and analyzed by western blotting. Antibodies used during Co-IP and subsequent IB include anti-MLL4 (anti-KMT2D/MLL4, Sigma # HPA035977), anti-P53 (Millipore, MABE327), and anti-IgG (Abcam, ab46540).

### Immunoblotting

Samples were separated by electrophoresis in 4%–20% SDS/PAGE gels with 20 mg per lane, transferred to PVDF membrane, and blotted with antibodies. Secondary horseradish peroxidase-conjugated secondary antibodies (Santa Cruz) and Amersham ECL Prime Western Blotting Detection Reagents (GE Healthcare, Cat# RPN2232) were used for detection. Antibodies used during immunoblotting include anti-actin (Cell Signaling, 4967S), anti-RARγ (Santa Cruz, sc-7387), anti-Alox12 (Santa Cruz, sc-365194).

### Histology

Mouse dorsal and ventral skin tissues were processed for histological examination by Core A of the Penn Skin Biology and Disease Resource-based Center and mounted on frost-free slides. All Hematoxylin and Eosin (H&E) staining was processed by the Penn Skin Biology and Disease Resource-based Center Core A. A Leica DM6 B microscope was used to observe and capture representative images. Exposure times and microscope intensity were kept constant for all human samples and across mouse littermate comparisons.

### Immunohistochemistry

Tissue slides were baked for one hour at 65°C, deparaffinized in xylene and rehydrated through a series of graded alcohols. After diH20 washes, slides were treated with Antigen Unmasking Solution (1:100, SKU H-3300-250, Vector Laboratories) at 95°C for 10 minutes according to the manufacturer’s protocol. Blocking and primary antibody binding was detected using the Vectastain Elite ABC-HRP system according to the manufacturer’s instructions (Cat#: PK-6200, Vector Laboratories). After O/N primary antibody (see antibody table) incubation at 4°C, endogenous peroxidases were blocked with 3% hydrogen peroxide in MeOH for 10 minutes at RT, treated with biotinylated secondary anti-mouse/rabbit antibody at RT for 90 min, and treated with ABC reagent (Vectastain Elite ABC-HRP Kit, PK-6200, Vector Laboratories) for 30 min at RT. The staining was visualized with 3,3’-diaminobenzidine (Cat#: SK-4100, Vector Laboratories) as peroxidase substrate. Exposure times were synchronized so that all tissues samples within an antibody group were exposed to DAB for the exact same time. All slides were counterstained with hematoxylin (Hematoxylin QS, H-3404, Vector Laboratories) for 35 seconds at RT, dehydrated in ethanol, cleared in xylene and mounted with VectaMount (Permanent mounting Medium, H-5000, Vector Laboratories). Antibodies used for IHC include anti-4-HNE (Abcam, ab46545), anti-Scnn1a (ProteinTech, 10924-2-AP), and anti-Alox12 (Santa Cruz, sc-365194).

### Immunofluorescence

Mouse tissues slides were deparaffinized with xylene, rehydrated with alcohol, and treated with Targeting Unmasking Fluid (1:3, Cat# Z000R.0000, Pan Path) at 90°C for 10 minutes according to the manufacturer’s protocol. Slides were permeabilized with 0.5% Triton X-100 in DPBS for 10 minutes and incubated in BlockAid Blocking Solution (Thermo Fisher, Cat# B10710) for 1 h at RT in a humidity chamber and incubated O/N in primary antibody. Following secondary antibody treatment (see antibody table) for 30 min at RT, the sections were mounted with ProLong Gold with DAPI (Thermo Fisher, Cat# P36935). Antibodies used for IF include anti-Krt14 (Abcam, ab7800), anti-Krt10 (Abcam, ab7638), anti-H3K4me1 (Abcam, ab8895), anti-RARγ (Santa Cruz, sc-7387), and anti-CD3 (Abcam, ab5690).

### Weight Collection

Following euthanasia, mice at 21 days of age were measured for total body weight. Data figures are the composite of multiple litters of Mll4-eKO or littermate controls. A one-way Anova followed by Tukey’s multiple comparison test was used to calculate significant differences between groups for all mice when stratified by gender and genotype. A student’s t-test was used to calculate significance between groups when considered for genotype only. All p values are noted in figure legends and were considered significant if p<0.05 and non-significant (NS) if p>0.05.

### TCGA analysis

RNA-seq data sets from head and neck squamous cell carcinoma and bladder cancer were obtained from TCGA (https://tcga-data.nci.nih.gov/tcga). Original RNA expression values (normalized read counts) were used for downstream analyses. For each cancer type, samples were ranked by MLL4 expression levels and were evenly divided into four groups. Comparisons were performed between the group of samples with the highest MLL4 expression and the group of samples with lowest MLL4 expression. One-sided Wilcoxon rank sum tests were used to compute significance.

### Human Protein Atlas

IHC data derived from the Human Protein Atlas can be found here: https://www.proteinatlas.org/

### Statistical Analyses

All statistical analyses were performed using R or Graphpad Prism 8. Details of each statistical test are included under the respective method section. Sample sizes and p-values are included in the figure legends or main figures. Investigators were not blinded during experiments or outcome assessment.

## DATA AND CODE AVAILABILITY

All data generated and code is available upon request, and all data will be deposited in GEO upon acceptance of the manuscript and prior to publication.

## ACKNOWLEDGEMENTS

This work was supported by the National Institute of Arthritis and Musculoskeletal and Skin Diseases (NIAMS) (K08AR070289 and R01 AR077615), the Damon Runyon Cancer Foundation, and the Dermatology Foundation, all to B.C.C. S.E. is supported by NIAMS (T32AR007465). We thank Steve Prouty for all of his assistance on tissue processing and histology.

## AUTHOR CONTRIBUTIONS

S.E., J.Z. and A.A. performed all experiments. S.E. and Y.A. performed all bioinformatic analysis. J.T.S. assisted with histological analyses. K.G. provided key reagents. S.E. and B.C.C. conceived of this work and wrote the manuscript. All authors have read and approved of the final manuscript.

## CONFLICTS OF INTEREST

The authors declare no competing interests.

---

**Extended Data Figure 1.**
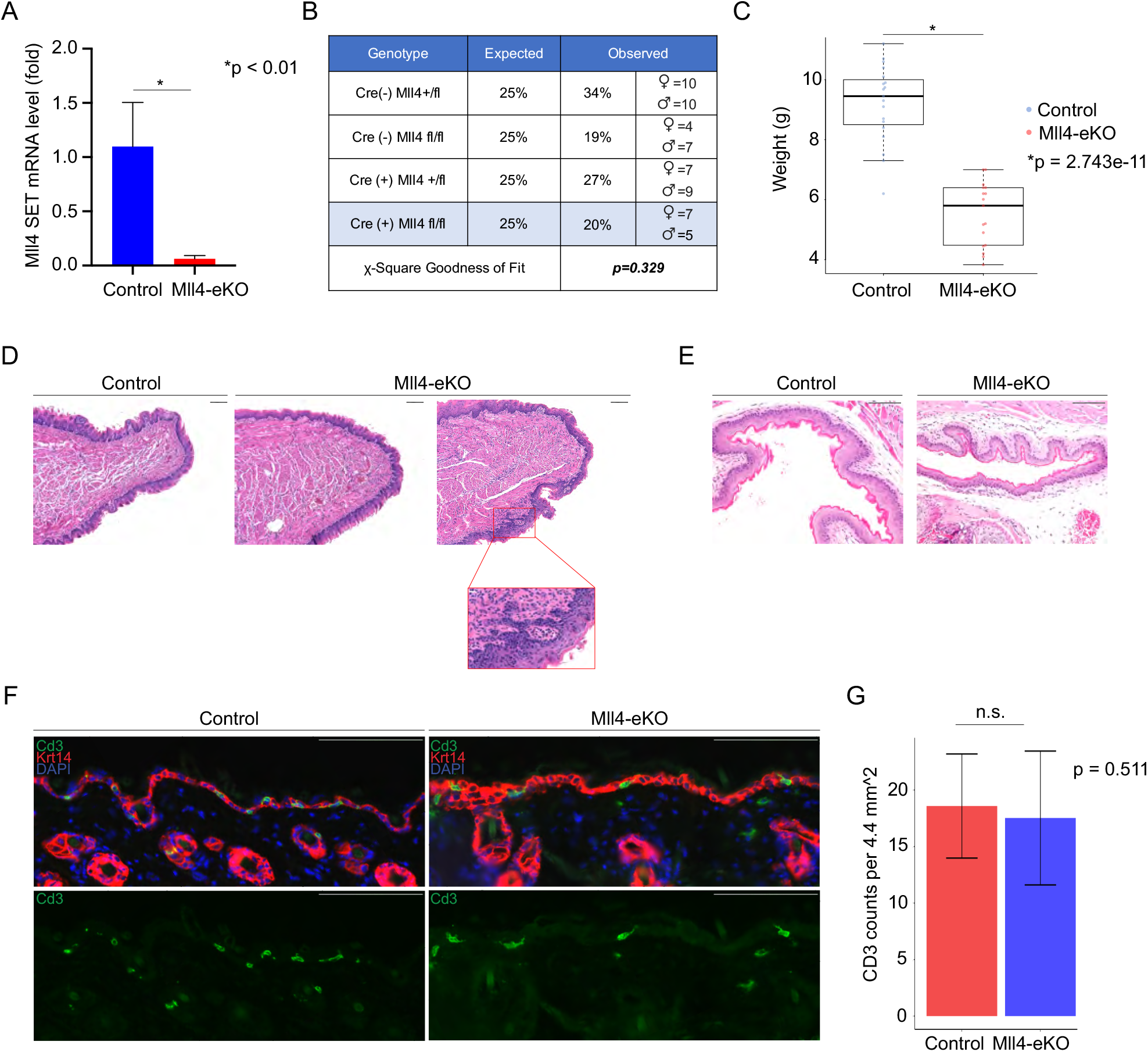
**(A)** qRT-PCR of Mll4-SET expression in isolated bulk epidermis of 3-week-old Mll4-eKO and control mice (n=3-4 mice per genotype, p<0.01). **(B)** Expected and observed percentiles of mendelian ratio birth rates of Mll4-eKO mice (p=0.329). **(C)** Weights of 3-week-old Mll4-eKO (red) and control (blue) mice (n=17-19 mice per genotype, p=2.743e-11)**. (D)** H&E histological staining of 3-week-old Mll4-eKO and control mice tongue (n=3 mice per genotype)**. (E)** H&E histological staining of 3-week-old Mll4-eKO and control mice esophagus (n=3 mice per genotype)**. (F)** IF staining of 3-week-old Mll4-eKO and control mice epidermis for Keratin-14 (red), CD3 (green), and DAPI (blue) (n=2 mice per genotype). **(G)** Quantification of total CD3 positive cells per 4.4mm^2 of images collected in F (p=0.511). Scale bar: 100 μM unless otherwise noted.

**Extended Data Figure 2.**
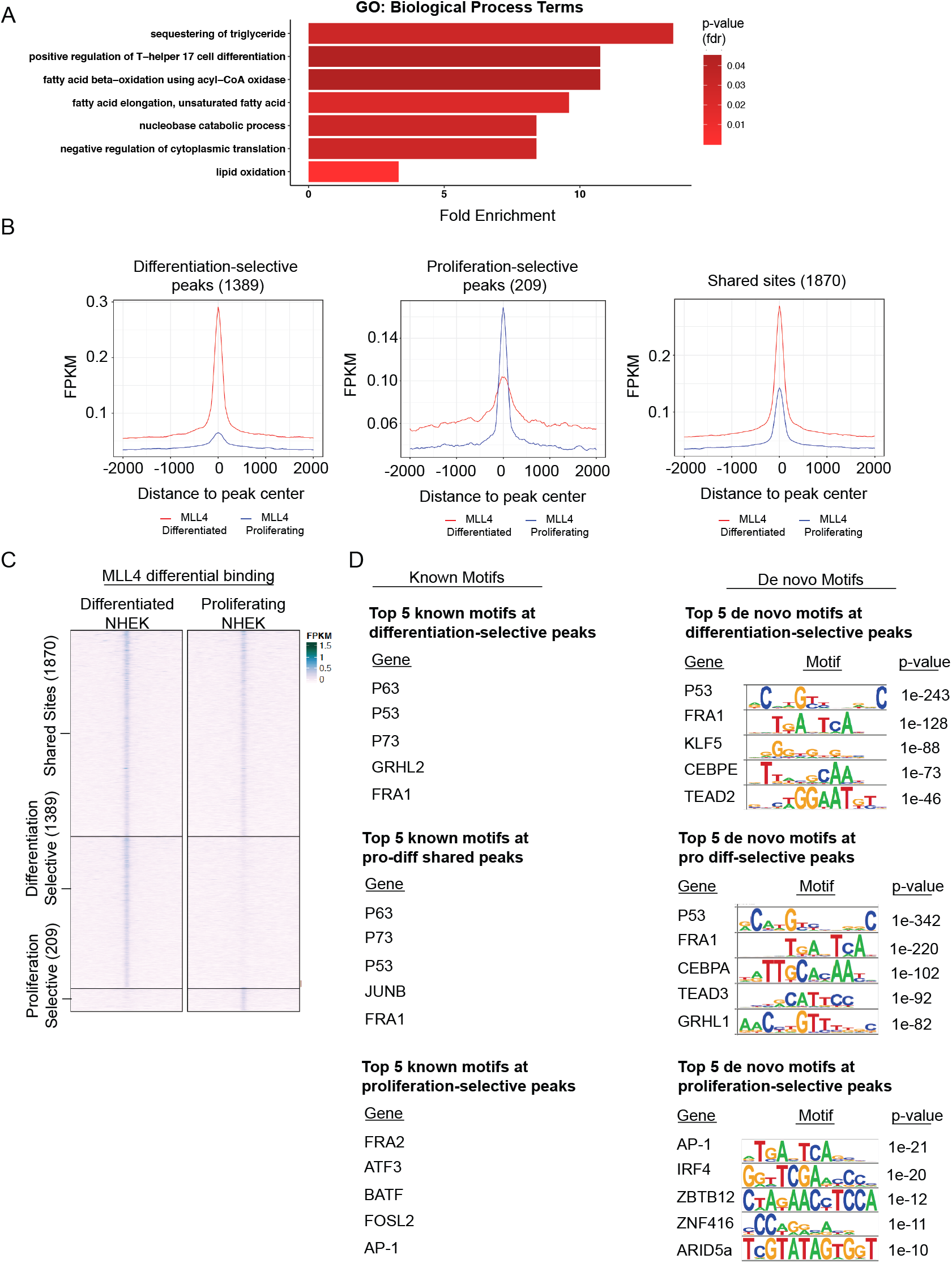
**(A)** qRT-PCR of Mll4-SET expression in isolated bulk epidermis of 3-week-old Mll4-eKO and control mice (n=3-4 mice per genotype, p<0.01). **(B)** Expected and observed percentiles of mendelian ratio birth rates of Mll4-eKO mice (p=0.329). **(C)** Weights of 3-week-old Mll4-eKO (red) and control (blue) mice (n=17-19 mice per genotype, p=2.743e-11). **(D)** H&E histological staining of 3-week-old Mll4-eKO and control mice tongue (n=3 mice per genotype). **(E)** H&E histological staining of 3-week-old Mll4-eKO and control mice esophagus (n=3 mice per genotype). **(F)** IF staining of 3-week-old Mll4-eKO and control mice epidermis for Keratin-14 (red), CD3 (green), and DAPI (blue) (n=2 mice per genotype). **(G)** Quantification of total CD3 positive cells per 4.4mm^2 of images collected in F (p=0.511). Scale bar: 100 μM unless otherwise noted.

**Extended Data Figure 3.**
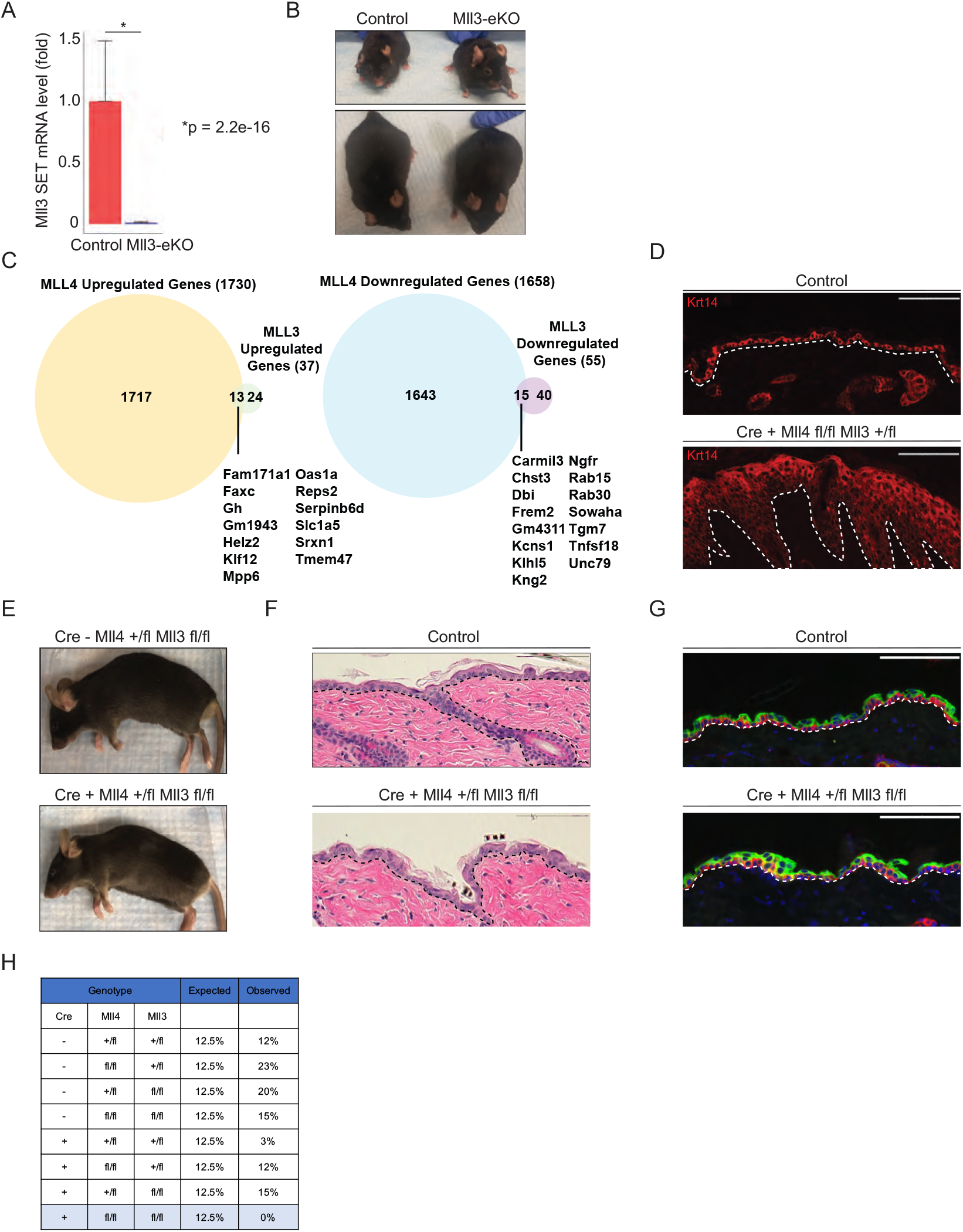
**A)** qRT-PCR of Mll3-SET expression in isolated bulk epidermis of 3-week-old Mll3-eKO and control mice (n=3-4 mice per genotype. p<0.01). **(B)** Representative 1–year-old Mll3-eKO and control mice. **(C)** Venn diagrams showing overlapping genes between Mll4-eKO and Mll3-eKO upregulated (left) or downregulated (right) genes. **(D)** IF staining of Krt14-Cre(+); *Mll4*^fl/fl^; *Mll3*^+/fl^ and control mice epidermis for Keratin-14 (red), and DAPI (blue) (n=3 mice per genotype). **(E)** Representative adult Krt14-Cre(+); *Mll4*^fl/+^; *Mll3*^fl/fl^ and control mice. **(F, H)** H&E histological staining of Krt14-Cre(+); *Mll4*^fl/+^; *Mll3*^fl/fl^ and control epidermis (n=3 mice per genotype). **(G)** IF staining of Krt14-Cre(+); *Mll4*^fl/+^; *Mll3*^fl/fl^ and control mice epidermis for Keratin-14 (red), Keratin-10 (green), and DAPI (blue) (n=3 mice per genotype). **(H)** Expected and observed percentiles for the offspring resultant from crosses between Krt14-Cre(+); *Mll4*^fl/+^; *Mll3*^fl/+^ and Krt14-Cre(−); *Mll4*^fl/fl^; *Mll3*^fl/fl^ mice (n=35 mice total). Scale bar: 100 μM unless otherwise noted.

## Notes

### Competing Interest Statement

The authors have declared no competing interest.

## REFERENCES

1 Bartha, I., di Iulio, J., Venter, J. C. & Telenti, A. Human gene essentiality. Nat Rev Genet 19, 51–62, doi:10.1038/nrg.2017.75 (2018).

2 Lee, J. E. et al. H3K4 mono- and di-methyltransferase MLL4 is required for enhancer activation during cell differentiation. Elife 2, e01503, doi:10.7554/eLife.01503 (2013).

3 Cerami, E. et al. The cBio cancer genomics portal: an open platform for exploring multidimensional cancer genomics data. Cancer Discov 2, 401–404, doi:10.1158/2159-8290.CD-12-0095 (2012).

4 Gao, J. et al. Integrative analysis of complex cancer genomics and clinical profiles using the cBioPortal. Sci Signal 6, pl1, doi:10.1126/scisignal.2004088 (2013).

5 Tate, J. G. et al. COSMIC: the Catalogue Of Somatic Mutations In Cancer. Nucleic Acids Res 47, D941–D947, doi:10.1093/nar/gky1015 (2019).

6 Fagan, R. J. & Dingwall, A. K. COMPASS Ascending: Emerging clues regarding the roles of MLL3/KMT2C and MLL2/KMT2D proteins in cancer. Cancer Lett 458, 56–65, doi:10.1016/j.canlet.2019.05.024 (2019).

7 Ford, D. J. & Dingwall, A. K. The cancer COMPASS: navigating the functions of MLL complexes in cancer. Cancer Genet 208, 178–191, doi:10.1016/j.cancergen.2015.01.005 (2015).

8 Jiang, X., Stockwell, B. R. & Conrad, M. Ferroptosis: mechanisms, biology and role in disease. Nat Rev Mol Cell Biol 22, 266–282, doi:10.1038/s41580-020-00324-8 (2021).

9 Stockwell, B. R., Jiang, X. & Gu, W. Emerging Mechanisms and Disease Relevance of Ferroptosis. Trends Cell Biol 30, 478–490, doi:10.1016/j.tcb.2020.02.009 (2020).

10 Chu, B. et al. ALOX12 is required for p53-mediated tumour suppression through a distinct ferroptosis pathway. Nat Cell Biol 21, 579–591, doi:10.1038/s41556-019-0305-6 (2019).

11 Hanahan, D. & Weinberg, R. A. Hallmarks of cancer: the next generation. Cell 144, 646–674, doi:10.1016/j.cell.2011.02.013 (2011).

12 Campbell, J. D. et al. Genomic, Pathway Network, and Immunologic Features Distinguishing Squamous Carcinomas. Cell Rep 23, 194–212 e196, doi:10.1016/j.celrep.2018.03.063 (2018).

13 Fowler, J. C. et al. Selection of Oncogenic Mutant Clones in Normal Human Skin Varies with Body Site. Cancer Discov 11, 340–361, doi:10.1158/2159-8290.CD-20-1092 (2021).

14 Lawson, A. R. J. et al. Extensive heterogeneity in somatic mutation and selection in the human bladder. Science 370, 75–82, doi:10.1126/science.aba8347 (2020).

15 Li, R. et al. Macroscopic somatic clonal expansion in morphologically normal human urothelium. Science 370, 82–89, doi:10.1126/science.aba7300 (2020).

16 Li, Y. Y. et al. Genomic analysis of metastatic cutaneous squamous cell carcinoma. Clin Cancer Res 21, 1447–1456, doi:10.1158/1078-0432.CCR-14-1773 (2015).

17 Pickering, C. R. et al. Mutational landscape of aggressive cutaneous squamous cell carcinoma. Clin Cancer Res 20, 6582–6592, doi:10.1158/1078-0432.CCR-14-1768 (2014).

18 Yilmaz, A. S. et al. Differential mutation frequencies in metastatic cutaneous squamous cell carcinomas versus primary tumors. Cancer 123, 1184–1193, doi:10.1002/cncr.30459 (2017).

19 Nehal, K. S. & Bichakjian, C. K. Update on Keratinocyte Carcinomas. N Engl J Med 379, 363–374, doi:10.1056/NEJMra1708701 (2018).

20 Lin-Shiao, E. et al. KMT2D regulates p63 target enhancers to coordinate epithelial homeostasis. Genes Dev 32, 181–193, doi:10.1101/gad.306241.117 (2018).

21 Jang, Y. et al. H3.3K4M destabilizes enhancer H3K4 methyltransferases MLL3/MLL4 and impairs adipose tissue development. Nucleic Acids Res 47, 607–620, doi:10.1093/nar/gky982 (2019).

22 Darido, C., Georgy, S. R. & Jane, S. M. The role of barrier genes in epidermal malignancy. Oncogene 35, 5705–5712, doi:10.1038/onc.2016.84 (2016).

23 Feingold, K. R. & Elias, P. M. Role of lipids in the formation and maintenance of the cutaneous permeability barrier. Biochim Biophys Acta 1841, 280–294, doi:10.1016/j.bbalip.2013.11.007 (2014).

24 Krieg, P. & Furstenberger, G. The role of lipoxygenases in epidermis. Biochim Biophys Acta 1841, 390–400, doi:10.1016/j.bbalip.2013.08.005 (2014).

25 Dang, N. & Murrell, D. F. Mutation analysis and characterization of COL7A1 mutations in dystrophic epidermolysis bullosa. Exp Dermatol 17, 553–568, doi:10.1111/j.1600-0625.2008.00723.x (2008).

26 Eckl, K. M. et al. Mutation spectrum and functional analysis of epidermis-type lipoxygenases in patients with autosomal recessive congenital ichthyosis. Hum Mutat 26, 351–361, doi:10.1002/humu.20236 (2005).

27 Fischer, J. et al. Mutations in the gene encoding SLURP-1 in Mal de Meleda. Hum Mol Genet 10, 875–880, doi:10.1093/hmg/10.8.875 (2001).

28 Nagarajan, S. R., Butler, L. M. & Hoy, A. J. The diversity and breadth of cancer cell fatty acid metabolism. Cancer Metab 9, 2, doi:10.1186/s40170-020-00237-2 (2021).

29 Wang, C. et al. Elevated level of mitochondrial reactive oxygen species via fatty acid beta-oxidation in cancer stem cells promotes cancer metastasis by inducing epithelial-mesenchymal transition. Stem Cell Res Ther 10, 175, doi:10.1186/s13287-019-1265-2 (2019).

30 Chen, M. & Huang, J. The expanded role of fatty acid metabolism in cancer: new aspects and targets. Precis Clin Med 2, 183–191, doi:10.1093/pcmedi/pbz017 (2019).

31 Stevens, B. M. et al. Fatty acid metabolism underlies venetoclax resistance in acute myeloid leukemia stem cells. Nat Cancer 1, 1176–1187, doi:10.1038/s43018-020-00126-z (2020).

32 Bergers, G. & Fendt, S. M. The metabolism of cancer cells during metastasis. Nat Rev Cancer 21, 162–180, doi:10.1038/s41568-020-00320-2 (2021).

33 Nomura, D. K. et al. Monoacylglycerol lipase regulates a fatty acid network that promotes cancer pathogenesis. Cell 140, 49–61, doi:10.1016/j.cell.2009.11.027 (2010).

34 Pascual, G. et al. Targeting metastasis-initiating cells through the fatty acid receptor CD36. Nature 541, 41–45, doi:10.1038/nature20791 (2017).

35 Charles, R. P. et al. Postnatal requirement of the epithelial sodium channel for maintenance of epidermal barrier function. J Biol Chem 283, 2622–2630, doi:10.1074/jbc.M708829200 (2008).

36 Szymanski, L. et al. Retinoic Acid and Its Derivatives in Skin. Cells 9, doi:10.3390/cells9122660 (2020).

37 Lee, J. et al. Targeted inactivation of MLL3 histone H3-Lys-4 methyltransferase activity in the mouse reveals vital roles for MLL3 in adipogenesis. Proc Natl Acad Sci U S A 105, 19229–19234, doi:10.1073/pnas.0810100105 (2008).

38 Lee, S. et al. Coactivator as a target gene specificity determinant for histone H3 lysine 4 methyltransferases. Proc Natl Acad Sci U S A 103, 15392–15397, doi:10.1073/pnas.0607313103 (2006).

39 Seiler, A. et al. Glutathione peroxidase 4 senses and translates oxidative stress into 12/15-lipoxygenase dependent- and AIF-mediated cell death. Cell Metab 8, 237–248, doi:10.1016/j.cmet.2008.07.005 (2008).

40 Ferretti, E., De Smaele, E., Di Marcotullio, L., Screpanti, I. & Gulino, A. Hedgehog checkpoints in medulloblastoma: the chromosome 17p deletion paradigm. Trends Mol Med 11, 537–545, doi:10.1016/j.molmed.2005.10.005 (2005).

41 Tam, C. S. & Stilgenbauer, S. How best to manage patients with chronic lymphocytic leuekmia with 17p deletion and/or TP53 mutation? Leuk Lymphoma 56, 587–593, doi:10.3109/10428194.2015.1011641 (2015).

42 Guo, Y. et al. Platelet 12-lipoxygenase Arg261Gln polymorphism: functional characterization and association with risk of esophageal squamous cell carcinoma in combination with COX-2 polymorphisms. Pharmacogenet Genomics 17, 197–205, doi:10.1097/FPC.0b013e328010bda1 (2007).

43 Yang, W. S. & Stockwell, B. R. Ferroptosis: Death by Lipid Peroxidation. Trends Cell Biol 26, 165–176, doi:10.1016/j.tcb.2015.10.014 (2016).

44 Savino, A. M. et al. Metabolic adaptation of acute lymphoblastic leukemia to the central nervous system microenvironment is dependent on Stearoyl CoA desaturase. Nat Cancer 1, 998–1009, doi:10.1038/s43018-020-00115-2 (2020).

45 Tesfay, L. et al. Stearoyl-CoA Desaturase 1 Protects Ovarian Cancer Cells from Ferroptotic Cell Death. Cancer Res 79, 5355–5366, doi:10.1158/0008-5472.CAN-19-0369 (2019).

46 Feng, H. & Stockwell, B. R. Unsolved mysteries: How does lipid peroxidation cause ferroptosis? PLoS Biol 16, e2006203, doi:10.1371/journal.pbio.2006203 (2018).

47 Uhlen, M. et al. Proteomics. Tissue-based map of the human proteome. Science 347, 1260419, doi:10.1126/science.1260419 (2015).

48 Soares, E. & Zhou, H. Master regulatory role of p63 in epidermal development and disease. Cell Mol Life Sci 75, 1179–1190, doi:10.1007/s00018-017-2701-z (2018).

49 Kim, S. et al. p63 directly induces expression of Alox12, a regulator of epidermal barrier formation. Exp Dermatol 18, 1016–1021, doi:10.1111/j.1600-0625.2009.00894.x (2009).

50 Kumar, R. et al. Mitochondrial uncoupling reveals a novel therapeutic opportunity for p53-defective cancers. Nat Commun 9, 3931, doi:10.1038/s41467-018-05805-1 (2018).

51 Wang, W. et al. CD8(+) T cells regulate tumour ferroptosis during cancer immunotherapy. Nature 569, 270–274, doi:10.1038/s41586-019-1170-y (2019).

52 Dobin, A. et al. STAR: ultrafast universal RNA-seq aligner. Bioinformatics 29, 15–21, doi:10.1093/bioinformatics/bts635 (2013).

53 Love, M. I., Huber, W. & Anders, S. Moderated estimation of fold change and dispersion for RNA-seq data with DESeq2. Genome Biol 15, 550, doi:10.1186/s13059-014-0550-8 (2014).

54 Egolf, S. et al. LSD1 Inhibition Promotes Epithelial Differentiation through Derepression of Fate-Determining Transcription Factors. Cell Rep 28, 1981–1992 e1987, doi:10.1016/j.celrep.2019.07.058 (2019).

55 Langmead, B. & Salzberg, S. L. Fast gapped-read alignment with Bowtie 2. Nat Methods 9, 357–359, doi:10.1038/nmeth.1923 (2012).

56 Li, H. et al. The Sequence Alignment/Map format and SAMtools. Bioinformatics 25, 2078–2079, doi:10.1093/bioinformatics/btp352 (2009).

57 Heinz, S. et al. Simple combinations of lineage-determining transcription factors prime cis-regulatory elements required for macrophage and B cell identities. Mol Cell 38, 576–589, doi:10.1016/j.molcel.2010.05.004 (2010).

58 Dou, Z. et al. Autophagy mediates degradation of nuclear lamina. Nature 527, 105–109, doi:10.1038/nature15548 (2015).

